# Learning spatio-temporal properties of hippocampal place cells

**DOI:** 10.1101/2021.07.13.452268

**Authors:** Yanbo Lian, Anthony N. Burkitt

## Abstract

Hippocampal place cells have spatio-temporal properties: they generally respond to a single spatial location of a small environment; in addition, they also display the temporal response property of theta phase precession, namely that the phase of spiking relative to the theta wave shifts from the late phase to early phase as the animal crosses the place field. Grid cells in layer II of the medial entorhinal cortex (MEC) also have spatio-temporal properties similar to hippocampal place cells, except that grid cells respond to multiple spatial locations that form a hexagonal pattern. Other non-grid spatial cells are also abundant in the entorhinal cortex (EC). Because the EC is the upstream area that projects strongly to the hippocampus, a number of EC-hippocampus models have been proposed to explain how the spatial receptive field properties of place cells emerge. However, none of these learning models have explained how the temporal response properties of hippocampal place cells emerge as a result of the EC input. A learning model is presented here based on non-negative sparse coding in which we show that the spatial and temporal properties of hippocampal place cells can be simultaneously learnt from EC input: both MEC grid cells and other EC spatial cells contribute to the spatial properties of hippocampal place cells while MEC grid cells predominantly determine the temporal response properties of hippocampal place cells.

## 1 Introduction

In early electrophysiological experiments involving freely behaving rats (O’Keefe and Dostrovsky, 1971; O’Keefe, 1976; Hill, 1978; O’Keefe and Conway, 1978), neuroscientists discovered place cells, the principal cells in the hippocampus. Place cells, as suggested by their name, have the spatial property of responding selectively to places in the external environment; namely they generally respond to a particular location (called the place field) of the spatial environment, although they may respond to multiple locations in a large environment (Park et al., 2011). In addition to the spatial properties of place cells, their temporal response property, namely theta phase precession, was observed by O’Keefe and Recce (1993): the place cell fires spikes at progressively earlier phases of the local field potential theta rhythm (7-12 Hz) while the animal moves across the place field. Normally, the firing of spike starts at the late phase of the theta rhythm when the animal enters the place field and ends at the early phase when the animal exits the place field (O’Keefe and Recce, 1993; Skaggs et al., 1996).

Three decades after hippocampal place cells were discovered, Hafting et al. (2005) reported another type of spatial cells, the grid cells, in the medial entorhinal cortex (MEC) that is an adjacent area to the hippocampus. Similar to the spatial properties of place cells in the hippocampus, MEC grid cells are also selective to spatial locations of the environment, but each MEC grid cell responds to multiple spatial locations (each location is called a grid field) forming a hexagonal grid that tiles the entire environment (Hafting et al., 2005). Moreover, there is a diversity resulting from multiple hexagonal grids of different MEC grid cells that have different orientations, spacings and offsets (Stensola et al., 2012). Subsequently, MEC grid cells were also observed to have the temporal response property of theta precession: the firing of MEC grid cells begins at the late phase of the theta rhythm when the animal enters the grid field, the phase of firing shifts in a systematic way during the traversal of the grid field, and ends at the early phase when the animal exits the grid field (Hafting et al., 2008). Though the spatial properties of place cells and MEC grid cells were discovered in an open environment (O’Keefe and Dostrovsky, 1971; Hafting et al., 2005), early studies primarily investigated theta precession on linear tracks (O’Keefe and Recce, 1993; Hafting et al., 2008). The phase precession of place cells and MEC grid cells in the open environment was investigated in later studies by Huxter et al. (2008), Climer et al. (2013) and Jeewajee et al. (2014).

Apart from grid cells in MEC, there are also many non-grid spatial cells in the MEC (Diehl et al., 2017; Hardcastle et al., 2017). In the lateral entorhinal cortex (LEC), many cells also contain spatial information (Hargreaves et al., 2005; Yoganarasimha et al., 2011). In this paper, we refer to these cells in the entorhinal cortex (EC) that contain spatial information but display no spatial structure like grid cells as EC weakly spatial cells.

Hippocampal place cells and MEC grid cells are fundamental units of the navigational system of the brain and there is a close relationship between these two types of cells. Experimental studies indicate that input from the entorhinal cortex (EC) is the principal input to the hippocampus (Steward and Scoville, 1976; Tamamaki and Nojyo, 1993; Leutgeb et al., 2007; Zhang et al., 2013). Consequently, MEC grid cells along with other cells in the EC are thought to provide spatial input for hippocampal place cells, so that they can have a specific place tuning. This has led to numerous feedforward EC-hippocampus models in which single place fields are generated from EC input, such as MEC grid cells or EC weakly spatial cells (Solstad et al., 2006; Rolls et al., 2006; Franzius et al., 2007a,b; Hasselmo, 2009; de Almeida et al., 2009; Savelli and Knierim, 2010; Neher et al., 2017; Lian and Burkitt, 2021). Furthermore, Fiete et al. (2008) showed that this MEC grid cell structure represents an encoding scheme for position that is analogous to the residue number system (Soderstrand et al., 1986), which is a highly efficient and accurate method for place representation (Sreenivasan and Fiete, 2011). On the other hand, place cells in CA1 of the hippocampus project back directly or indirectly to the EC (Kloosterman et al., 2003; Naber et al., 2001; Slomianka et al., 2011) and the inactivation of place cells leads to the degradation of receptive field structure of MEC grid cells (Bonnevie et al., 2013), supporting a loop-like EC-hippocampus network structure. This loop-like structure has been adopted in some modelling studies to explain other spatial properties of place and MEC grid cells such as global and rate remapping of place cells, and multisensory integration in MEC grid cells (Rennó-Costa and Tort, 2017; Li et al., 2020). Compared with the feedforward EC-hippocampus models, the loop-like models can also incorporate the effects on the spatial firing of MEC grid cells due to the feedback introduced in such models.

Separate from the link between the spatial properties of place cells and MEC grid cells, some experimental studies also infer a link between the temporal properties of place cells and MEC grid cells. Hafting et al. (2008) discovered that MEC grid cells have hippocampus-independent phase precession and suggested that phase precession of hippocampal place cells could be inherited from the EC. Yamaguchi et al. (2007) proposed a entorhinal-hippocampal network that hypothesises that theta phase precession originates at EC superficial layers and is transmitted to the hippocampus along the hippocampal trisynaptic circuit. In a review paper, Bush et al. (2014) proposed that theta phase precession of place cells is likely inherited from MEC grid cell input. In a later study by Schlesiger et al. (2015), the MEC was found to be necessary for the temporal properties of hippocampal neuronal activity, even when place cells maintain stable spatial firing. However, existing EC-hippocampus models only focus on the learning of spatial properties and there is no existing model that explicitly shows how the temporal properties of hippocampal place cells can be inherited from MEC grid cells through some form of Hebbian learning. We propose here a learning model in which the spatio-temporal properties of hippocampal place cells emerge through synaptic plasticity during navigation of a virtual rat in a 2D environment.

The learning model presented here is built upon our previous work that shows that the model based on non-negative sparse coding can learn an efficient hippocampal place map using EC input such as MEC grid cells and EC weakly spatial cells (Lian and Burkitt, 2021). In this paper, building upon the work of Jeewajee et al. (2014) and McClain et al. (2019), we construct a mathematical model of MEC spatio-temporal grid cells of a 2D environment. These MEC spatio-temporal grid cells, along with EC weakly spatial cells, form the input to modelled hippocampal cells, and the connection between EC input cells (MEC grid cells and EC weakly spatial cells) and modelled hippocampal cells are learnt during navigation of a virtual rat. After learning, the modelled hippocampal cells display spatial and temporal properties that are similar to experimental data of place cells. Furthermore, after the learning process, these learnt hippocampal place cells still maintain their spatial selectivity even when MEC spatio-temporal grid cells are inactivated, suggesting that the remaining EC weakly spatial cells can maintain the place field responsiveness of place cells. Combining our previous work showing that either MEC grid cells or EC weakly spatial cells can give rise to the spatial tuning of hippocampal place cells (Lian and Burkitt, 2021), we complete the picture of how EC input contributes to the properties of hippocampal place cells from the perspective of synaptic plasticity; namely that hippocampal place cells learn the spatial properties from both MEC grid cells and EC weakly spatial cells, and learn their temporal properties from MEC grid cells. The synaptic plasticity underlying the learning model here is based on the principle of sparse coding.

## 2 Methods

### 2.1 The environment and trajectory

The spatial environment used in this study is a 1m×1m square box where a virtual rat runs freely. Similar to the study by D’Albis and Kempter (2017), the running trajectory ***r***_*t*_ is generated from the stochastic process described as

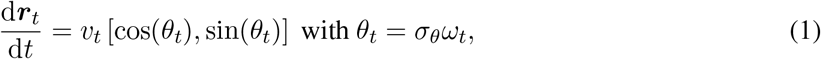

where *v*_*t*_ is the speed sampled from an Ornstein-Uhlenbeck process with long-term mean 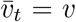, *θ*_*t*_ is the direction of movement, *ω*_*t*_ is a standard Wiener process, and *σ*_*θ*_ is the parameter that controls the tortuosity of the running trajectory.

When the virtual rat is running toward the wall and very close to the wall (within 2 cm), the running direction of the rat (*θ*_*t*_) is set to the direction parallel to the wall. If the rat location generated by Eq. 1 falls outside of the environment, the stochastic process keeps running until a valid location is generated.

The running trajectory of the virtual rat is generated at 100 Hz; i.e., the position is updated every 10 ms according to Eq. 1. The long-term mean speed, *v*, is set to 30 cm/s. A summary of parameters and values used in the simulation can be found in Table 1.

**Table 1:** Model symbols and parameters.

| Description (value) | Symbol |
| --- | --- |
| Tortuosity of the running trajectory (1 radian) | $\sigma_\theta$ |
| Spatial location / running speed / running direction and at time $t$ | $\mathbf{r}_t / v_t / \theta_t$ |
| Long-term mean of the speed (30 cm/s) | $v$ |
| Time step of the running trajectory (10 ms) |  |
| Duration of the training / testing running trajectory (3600 s / 1200 s) |  |
| Spatial location in the environment | $\mathbf{r}$ |
| Spatio-temporal response at spatial location $\mathbf{r}$ at time $t$ | $f(\mathbf{r}, t)$ |
| Spatial receptive field | $f_s(\mathbf{r})$ |
| Amplitude / centre / radius of the spatial receptive field | $\alpha_c / \mathbf{r}_c / R$ |
| Phase modulation at spatial location $\mathbf{r}$ at time $t$ | $f_\phi(\mathbf{r}, t)$ |
| Level of phase modulation (uniformly sample from [0.8 1.2]) | $k_\phi$ |
| Frequency of the theta rhythm (10 Hz) | $F$ |
| Firing phase at location $\mathbf{r}$ | $\phi(\mathbf{r})$ |
| Entry phase of phase precession (uniformly sampled from [300° 340°]) | $\phi_0$ |
| Phase change of phase precession (uniformly sampled from [300° 340°]) | $\Delta\phi$ |
| Projected centre / radius of the spatial receptive field onto running direction | $\mathbf{r}' / R'$ |
| Fitted spatial receptive fields of modelled hippocampal cells | $\hat{f}_s(\mathbf{r})$ |
| Amplitude / centre / radius of the fitted spatial receptive field | $\hat{\alpha}_c / \hat{\mathbf{r}}_c / \hat{R}$ |
| Fitted level of phase modulation | $\hat{k}_\phi$ |
| Fitted frequency of theta rhythm | $\hat{F}$ |
| Fitted phase of the temporal response | $\hat{\phi}$ |
| Membrane time constant of modelled hippocampal cells (10 ms) | $\tau$ |
| Membrane potentials of modelled hippocampal cells | $\mathbf{u}$ |
| Firing rates of modelled hippocampal cells | $\mathbf{s}$ |
| Firing rates of EC input cells | $\mathbf{I}$ |
| Excitatory connection: EC input cells to modelled hippocampal cells | $\mathbf{A}$ |
| Firing threshold of modelled hippocampal cells (0.3) | $\beta$ |
| Learning rate (0.01) | $\eta$ |
| Time step of computing modelled hippocampal cells by Eq. 7 (0.2 ms) |  |
| Number of modelled hippocampal cells in Scenario 1, 2 and 3 (100) |  |
| Number of MEC spatio-temporal grid cells in Scenario 1, 2 and 3 (900) |  |
| Number of EC weakly spatial cells in Scenario 2 and 3 (400) |  |

### 2.2 Model of MEC spatio-temporal grid cells

MEC grid cells are found to have spatial and temporal properties, namely they are selective to a hexagonal grid of spatial locations (Hafting et al., 2005) and have diverse grid parameters (Stensola et al., 2012), and their firing displays theta phase precession (Hafting et al., 2008). The spatial receptive fields of MEC grid cells can be characterised by the sum of three sinusoidal gratings (Solstad et al., 2006) or the sum of circular fields at different locations in the environment. In order to investigate the question of whether the spatial and temporal properties of hippocampal place cells can be inherited from MEC grid cells via learning, we first need a mathematical model that can characterise both spatial and temporal properties of MEC grid cells.

Burgess (2008) designed a model of grid cells based on the oscillatory interference model (O’Keefe and Burgess, 2005; Burgess et al., 2007). This model of grid cells consists of multiple membrane potential oscillators whose frequencies linearly depend on the speed and can capture the spatial and temporal properties of the grid cells. However, a recent experimental study challenged the classical view that the oscillation frequency linearly depends on the running speed (Kropff et al., 2021). Instead, the frequency of the theta rhythm is controlled by the acceleration (Kropff et al., 2021). In this study we build a mathematical model of spatio-temporal grid cells from a different perspective, inspired by the work of Chadwick et al. (2015) and McClain et al. (2019), in which the spatio-temporal responses of grid cells are modelled as the product of spatial and temporal responses as described below.

Chadwick et al. (2015) and McClain et al. (2019) build a mathematical model for 1D place cells that have spatial and temporal properties, and Jeewajee et al. (2014) showed that theta phase precession of grid cells and place cells strongly depends on the projected distance on the current running direction (pdcd) in a 2D environment. Analogous to their work, we construct a mathematical model of MEC spatio-temporal grid cells based on 2D spatio-temporal place cells. The fundamental idea is that the receptive field of each MEC spatio-temporal grid cell is seen as the combination of multiple grid fields located in the hexagonal grid where each grid field has spatio-temporal properties similar to a 2D spatio-temporal place cell.

Within the region of any receptive field, the firing rate at location ***r*** and time *t* is modelled as the product of a spatial receptive field and a phase modulation (McClain et al., 2019):

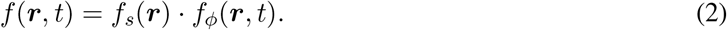

The spatial receptive field, *f*_*s*_(***r***), is described as a 2D function

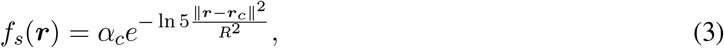

where ***r***_*c*_ is the centre of the receptive field, *α*_*c*_ is the amplitude at the centre, and *R* determines the radius of the spatial receptive field. Similar to our previous work (Lian and Burkitt, 2021), the spatial receptive field of each MEC spatio-temporal grid cell is determined by randomly sampling the grid spacing, orientation and offset from some distributions. Stensola et al. (2012) showed that MEC grid cells are discretised into modules based on their grid spacings. Following the model of Neher et al. (2017), we take four discrete grid modules whose mean grid spacings are 38.8 cm, 48.4 cm, 65 cm and 98.4 cm, and mean grid orientations are 15°, 30°, 45° and 0°, respectively. Similar to the distribution of grid spacings in discrete grid modules (Stensola et al., 2012), these four discrete grid modules account for 43.5%, 43.5%, 6.5% and 6.5% of grid cells in the model, respectively. For any MEC grid cell in its module, the grid spacing is sampled from a normal distribution with mean spacing and standard deviation of 8 cm, the grid orientation is sampled from a normal distribution with mean orientation and standard deviation of 3°, and the grid offset is sampled from a uniform distribution between 0 and the grid spacing. Because Ismakov et al. (2017) observed the variability in individual grid fields of a MEC grid cell, the amplitude (*α*_*c*_ in Eq. 3) of each grid field for a MEC grid cell is sampled from a normal distribution with mean 1 and standard deviation 0.1 (Neher et al., 2017; Lian and Burkitt, 2021). The radius, *R* in Eq. 3, of the receptive field is determined by *R* = 0.32*λ*, where *λ* is the grid spacing of MEC grid cell. The spatial receptive field (rate map) of one example MEC grid cell is displayed in Fig. 1A.

**Figure 1:**
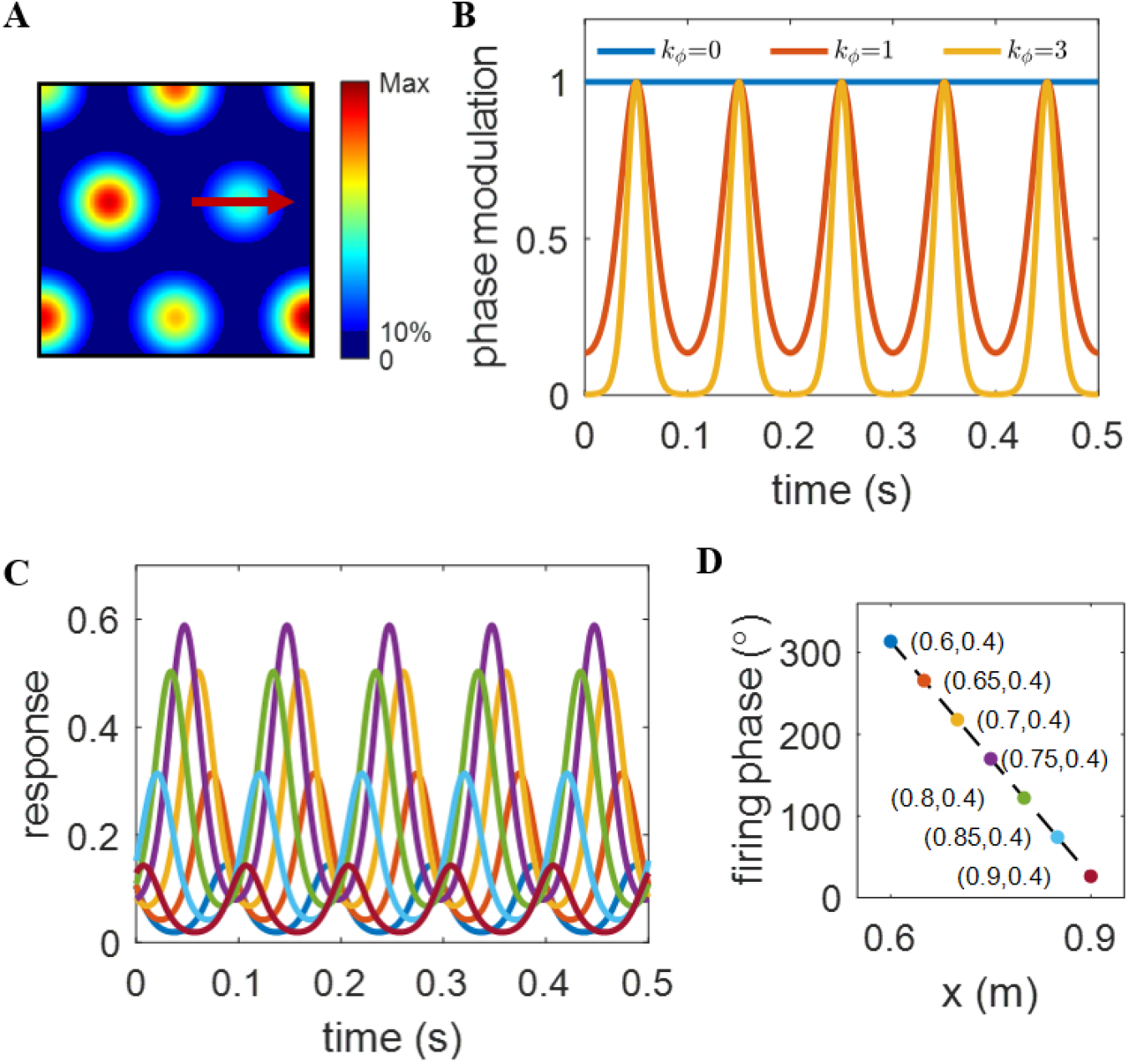
Illustration of the mathematical model of a MEC spatio-temporal grid cell. For this cell, *k*_*ϕ*_ = 1, *ϕ*_0_ = 320°, Δ*ϕ* = 300°, *R* = 0.16 m and *F* = 10 Hz. (A) The spatial receptive field (rate map), *f*_*s*_(***r***), in Eq. 3, of a MEC grid cell. (B) Phase modulation, *f*_*ϕ*_ (***r***, *t*), in Eq. 4, for 3 different values of *k*_*ϕ*_. *ϕ*(***r***) is set to 180°. The response is more phase locked for larger values of *k*_*ϕ*_. (C) Curves in different colours represent responses, *f* (***r***, *t*), of one pre-built spatio-temporal grid cell over 0.5 s at different locations (coordinates found in plot D with the same colour coding) when the virtual rat moves from left to right of the place field along the trajectory in plot A. (D) Firing phase, *ϕ*(***r***), vs. position. Only *x* varies because the animal moves straight from left to right.

The phase modulation, *f*_*ϕ*_(***r***, *t*), describes the temporal response (i.e., the probability of the timing of the individual action potentials) and is given by

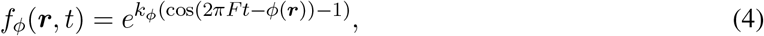

where *k*_*ϕ*_ is the parameter that controls the level of phase modulation and hence the extent of the resulting phase precession, *F* is the frequency of theta rhythm and set to 10 Hz throughout the paper. The choice of *F* = 10 Hz help with visualizing the results as every 0.1 s is a full period (see examples in Fig. 1B & C) and other choices of *F* will not change the results, apart from the frequency of the theta rhythm. *ϕ*(***r***) is a function that returns the firing phase at location ***r***. Fig. 1B shows the dependence of phase modulation on *k*_*ϕ*_. *ϕ*(***r***) is set to 180° when plotting the curve. The larger *k*_*ϕ*_ is, the stronger the phase modulation is. When *k*_*ϕ*_ is 0, there is no phase modulation and the response of MEC grid cells only depends on the spatial location of the animal.

In a 1D linear track, Chadwick et al. (2015) and McClain et al. (2019) model *ϕ*(*r*) as a linear function of the location, *r*:

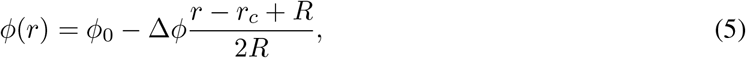

where *ϕ*_0_ is the entry phase at location *r*_*c*_ − *R*, Δ*ϕ* is the phase change across the receptive field, and thus the exit phase at location *r*_*c*_ + *R* is *ϕ*_0_ − Δ*ϕ*. Note that in 1D *r* and *r*_*c*_ become scalars instead of vectors. An example of the linear relationship between *ϕ*(*r*) and *r* described in Eq. 5 is shown in Fig. 1D.

However, determining *ϕ*(***r***) is more complicated in a 2D environment because the animal can enter and exit the receptive field from any location with any running direction, and the running trajectory does not always pass through the centre of the receptive field. Jeewajee et al. (2014) show that the best correlate of phase precession in a 2D open environment is the projected distance onto the animal’s current running direction (pdcd). Therefore, we model *ϕ*(***r***) in a 2D environment by a projected version of *ϕ*(*r*) in Eq. 5, described by:

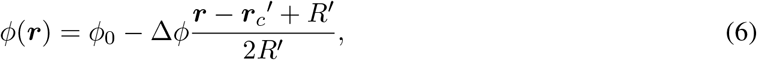

where 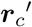 is the projected centre and *R*′ is the projected radius onto the running direction. In any location within the receptive field, the firing phase *ϕ*(***r***) depends on both the spatial location and running direction. Firing phases at the same location of different trajectories could have different values due to different running directions.

Above all, the spatio-temporal responses of a MEC grid cells in a 2D environment are simply modelled as multiple grid fields at all the vertices of the hexagonal grid in which each grid field is characterised by the spatio-temporal model described above (Eqs. 2-4 and 6).

From Eq. 4, we can see that the spatio-temporal response is periodical with frequency *F* when the virtual rat stays at a fixed location within the receptive field. Therefore, supposing the virtual rat remains stationary at ***r*** for 1 s, the spatio-temporal response over this 1 s will oscillate at frequency *F* with phase determined by *ϕ*(***r***) and the magnitude of the oscillating response is determined by *f*_*s*_(***r***). In other words, from the spatio-temporal response over 1 s at a fixed location, we can infer the firing phase and firing frequency of the temporal coding at the location. Though theta waves are generally absent in stationary animals (Bland, 1986), the spatio-temporal firing response over 1 s at a fixed location is used in this paper to visualise, estimate and analyse the temporal properties at different locations within the receptive field. However, this does not imply that the virtual rat stays at any location for 1 s when freely exploring the environment. When the model is in the learning stage, the virtual rat is continually moving in a spatial environment along a trajectory generated by Eq. 1. After the learning stage, the spatio-temporal response over 1 s at different locations is used to recover the learnt temporal properties of hippocampal place cells (details found in Section 2.8 and 2.9).

The response of one example MEC spatio-temporal grid cell is illustrated in Fig. 1. The spatial receptive field (rate map) of this grid cell is displayed in Fig. 1A. For this cell, *k*_*ϕ*_ = 1, *ϕ*_0_ = 320°, Δ*ϕ* = 300° and *R* = 0.16 m. The red arrow in Fig. 1A indicates the straight trajectory the virtual rat runs through a grid field from left to right. Fig. 1C shows the response over 0.5 s when the virtual rat is at different locations of the running trajectory (red arrow in Fig. 1A) in which the coordinates of locations are given in Fig. 1D with the same colour coding. At each location, the response over 0.5 s is intentionally plotted to visualise the spatial and temporal properties. As the virtual rat runs from the left to the right of the grid field, the spatial response (*f*_*s*_(***r***) in Eq. 3), describing the amplitude of the periodic curve, increases, peaks at the centre, and then decreases. However, the phases of these periodic curves (Fig. 1C) start with a value close to 360° and then keep decreasing along the trajectory, mimicking theta phase precession. Theta phase precession can also be observed from Fig. 1D which shows that the curve of firing phase vs. position (*x*) has a correlation coefficient −1. Note that theta phase precession is incorporated into the model through the temporal way in which the firing phase varies linearly with the pdcd within the receptive field (Eqs. 4 and 6). In this way Fig. 1 illustrates that our mathematical model of MEC spatio-temporal grid cells shows both spatial and temporal dependence, and the theta phase precession introduced here is perfect with a correlation coefficient −1. Naturally, the theta phase precession is unlikely to be so perfect in experimental studies. However, our aim here is to investigate from a modelling perspective how well the modelled hippocampal place cells preserve the theta phase precession of MEC spatio-temporal grid cells after learning.

### 2.3 EC weakly spatial cells

Though MEC grid cells have a clear spatial structure, they account for only approximately 20% of the MEC cell population (Diehl et al., 2017). Furthermore, Diehl et al. (2017) found that about two-thirds of MEC cells (i.e., the non-grid spatial cells) have less specialised but consistent spatial firing patterns. Hardcastle et al. (2017) discovered that many non-grid cells in the MEC have firing patterns that contain spatial information. In addition, cells in LEC, that also contain spatial information (Hargreaves et al., 2005; Yoganarasimha et al., 2011), can likewise contribute to the formation of hippocampal place cells. In our previous work, we have shown that a model based on sparse coding can effectively retrieve place information from EC weakly spatial cells that have no structured spatial selectivity to form an efficient hippocampal place map (Lian and Burkitt, 2021). In this paper, we incorporated EC weakly spatial cells and treated them as potential upstream input to the hippocampus to investigate how EC weakly spatial cells and MEC spatio-temporal grid cells altogether contribute to the spatial and temporal properties of modelled hippocampal place cells. In this model, the firing field of EC weakly spatial cells is generated by first sampling from a uniform distribution between 0 and 1 for each location, then smoothing the map with a Gaussian kernel with a standard deviation of 6 cm, and normalising the map to give values between 0 and 0.1.

### 2.4 Non-negative sparse coding

Similar to our previous work (Lian and Burkitt, 2021), we build here the learning model of spatio-temporal place cells based upon non-negative sparse coding (Hoyer, 2003). Sparse coding, originally proposed by Olshausen and Field (1996, 1997), finds an efficient representation of the input using a linear combination of some basis vectors. Non-negative sparse coding constrains the responses and basis vectors to be non-negative. We implement the model via a locally competitive algorithm (Rozell et al., 2008) that efficiently solves sparse coding as follows:

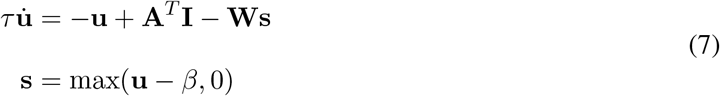

and

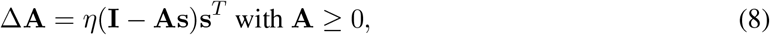

where **I** is the input, **s** represents the response (firing rate) of the model units, **u** can be interpreted as the membrane potential, **A** is the matrix containing basis vectors and can be interpreted as the connection weights between the input and model units, **W** = **A**^*T*^ **A** − 𝟙 and can be interpreted as the recurrent connection between model units, 𝟙 is the identity matrix, *τ* is the time constant, *β* is the positive sparsity constant that controls the threshold of firing, and *η* is the learning rate. Each column of **A** is normalised to have length 1. The non-negative constraints are incorporated into the system, as seen from the non-negativity of both **s** and **A** in Eqs. 7 and 8. The parameters of implementing Eqs. 7 and 8 are given below (Section 2.6). Additional details about the implementation of non-negative sparse coding can be found in Lian and Burkitt (2021).

### 2.5 Structure of the model

In this paper, the EC provides upstream input for the hippocampus and the EC-hippocampus pathway is modelled in three scenarios. Modelled hippocampal cells undergoes a learning process described by Eqs. 7 and 8. A modelled hippocampal cell is referred to as ‘learnt hippocampal place cell’ if it meets the criteria of a place cell after learning (Section 2.7.2). The diagram of the model structure is displayed in Fig. 2. A summary of all symbols defined in this paper is shown in Table 1.

**Figure 2:**
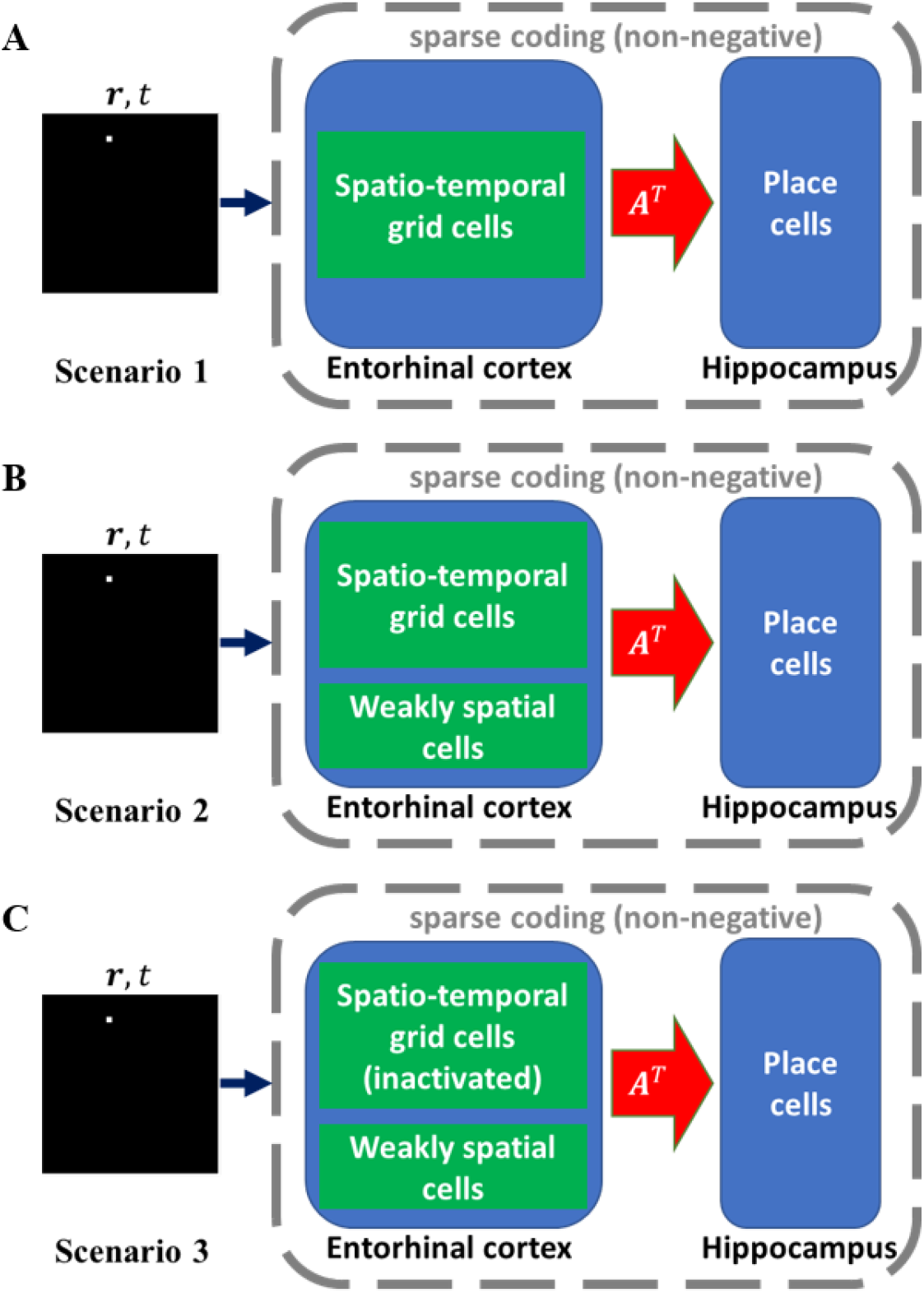
Model structure in three scenarios. MEC spatio-temporal grid cells or EC weakly spatial cells serve as the upstream input to the hippocampus. Given a spatial location ***r*** from a virtual rat running trajectory and time *t*, EC provides input for the hippocampus. The response of modelled hippocampal cells and the connection weights **A** are updated as described by Eqs. 7 and 8. (A) Scenario 1: only MEC spatio- temporal gird cells serve as the EC input. (B) Scenario 2: both MEC spatio-temporal grid cells and EC weakly spatial cells serve as the EC input. (C) Scenario 3: MEC spatio-temporal grid cells were inactivated after the learning process of Scenario 2.

**Scenario 1:** modelled hippocampal cells only receive input from MEC spatio-temporal grid cells, as described above, which have both spatial and temporal response properties (see Section 2.2). The EC-hippocampus connection undergoes a learning process while the virtual rat is exploring the environment. This scenario is designed to investigate the contribution of MEC spatio-temporal grid cells to the spatial and temporal properties of learnt hippocampal place cells.

**Scenario 2:** modelled hippocampal cells receive input from MEC spatio-temporal grid cells together with EC weakly spatial cells. Similar to Scenario 1, the EC-hippocampus connection is learnt. This scenario aims to validate and extend the results of Scenario 1 to the situation in which other spatial cells in the EC also contribute to the firing of learnt hippocampal place cells.

**Scenario 3:** after the learning process of Scenario 2, MEC spatio-temporal grid cells are inactivated and there is no learning in this scenario. This scenario is used to replicate the experimental setup of the study by Schlesiger et al. (2015), in which rats received NMDA lesions of the MEC. Schlesiger et al. (2015) found that theta phase precession of hippocampal place cells was greatly disrupted by MEC lesions but hippocampal spatial firing still remained. This scenario is also similar to another experimental study that showed the retention of hippocampal place fields during medial septum inactivation (Brandon et al., 2014), which is presumed to shut off grid cell input (Koenig et al., 2011; Brandon et al., 2011). In this scenario, the spatial firing of learnt hippocampal place cells will be investigated after the inactivation of MEC spatio-temporal grid cells in order to compare the simulation results of our model with the results of these experimental studies.

### 2.6 Training

The values of training parameters and definition of symbols can be found in Table 1. As the virtual rat moves in the environment, each MEC spatio-temporal grid cell generates a firing rate that is determined by the spatial location ***r*** and time *t* according to Eqs. 2-4 and 6, while each of the EC weakly spatial cells generates a firing rate determined only by the spatial location. The response of modelled hippocampal cells is computed iteratively using the model dynamics (Eq. 7). The connection, **A**, between EC cells and modelled hippocampal cells is updated according to the learning rule described in Eq. 8. There are 100 modelled hippocampal cells in the model and the three different scenarios above are implemented.

**Scenario 1:** only MEC spatio-temporal grid cells provide input to the modelled hippocampal cells, there are 900 MEC spatio-temporal grid cells; i.e., **A** is a 900 × 100 matrix. A running trajectory of 3600 s is used to train the model and the connection weight matrix, **A**, is learnt during this training process.

**Scenario 2:** both MEC spatio-temporal grid cells and EC weakly spatial cells provide input to the hippocampus. In addition to the 900 MEC spatio-temporal grid cells, there are also 400 EC weakly spatial cells; i.e., **A** is a 1300 × 100 matrix where the top 400 rows of **A** represent the connection weights from EC weakly spatial cells to modelled hippocampal cells and the bottom 900 rows represent the connection weights from MEC grid cells to modelled hippocampal cells. The same running trajectory of 3600 s, as used in Scenario 1, is used to train the model.

**Scenario 3:** after the learning process in Scenario 2, MEC spatio-temporal grid cells were inactivated in order to investigate how this affects the spatial firing of learnt hippocampal place cells after spatio-temporal input from MEC grid cells is lost. MEC grid cells are inactivated by setting the bottom 900 rows of **A** (the connection weights from MEC spatio-temporal grid cells to modelled hippocampal cells) to zero and then normalising each column of **A** to have length 1. In this scenario there is no learning.

The connection weight matrix **A** is initialised according to a uniform distribution between 0 and 1 and then normalised to have length 1 for each column. For both Scenarios 1 and 2, a running trajectory of 3600 s is used to train the model and the connection weight matrix, **A**, is learnt during this training process. After learning, another running trajectory of 1200 s is used to recover the spatial receptive fields of these 100 modelled hippocampal cells for all three scenarios.

The parameters in Eqs. 4 and 6 that control the temporal properties of MEC grid cells are chosen as follows: *k*_*ϕ*_ is chosen randomly from a uniform distribution between 0.8 and 1.2; *ϕ*_0_ and Δ*ϕ* are both chosen randomly from a uniform distribution between 300° and 340°; and *F* is 10 Hz.

For the parameters in the model dynamics and learning rule (Eqs. 7 and 8), *τ* is 10 ms, *β* is 0.3, and the time step for simulating the modelled hippocampal cells by Eq. 7 is taken to be 0.2 ms. Because the trajectory is updated after every 10 ms, there are 50 iterations of computing the response of modelled hippocampal cells using Eq. 7. The connection weight matrix, **A**, is updated using Eq. 8 after 50 iterations and the learning rate *η* is 0.01.

### 2.7 Recovering the spatial receptive fields of modelled hippocampal cells

After training, another running trajectory of the virtual rat with a duration of 1200 s was used to recover the spatial receptive fields (rate maps) of modelled hippocampal cells. The 1m×1m environment is discretised into a 40×40 lattice and the receptive field is recovered as the averaged response across all locations along the running trajectory of 1200 s.

#### 2.7.1 Fitting the spatial receptive fields

In order to quantitatively characterise the spatial receptive fields of learnt hippocampal cells, the recovered receptive fields using the approach above is fitted by the following function:

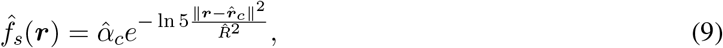

where 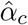 is the amplitude, 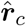 is the centre and 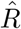 determines the radius of the receptive field. The fitting routine is performed using the built-in MATLAB (version R2020a) function *lsqcurvefit* and the fitting error is defined as the ratio between the summed square of the fitting residual and the summed square of the recovered receptive field.

#### 2.7.2 Selecting place cells

The criteria for categorising a modelled hippocampal cell as a place cell are: (1) The centre, 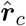, is inside the spatial environment, (2) The radius, 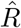, is larger than 5 cm, and (3) The fitting error is smaller than 40%, where the fitting error is defined as the square of the ratio between the fitting residual and spatial field. The receptive field of a place cell is called the *place field*. These last two criteria are designed to preserve learnt hippocampal place cells that have a reasonable size of the place field and only one dominant place field. A modelled hippocampal cell that meets the criteria is referred to as ‘learnt hippocampal place cell’.

### 2.8 Fitting the spatio-temporal response of learnt hippocampal place cells

The MEC spatio-temporal grid cells in the model have built-in spatial and temporal properties as described in Eqs. 2-4 and 6, so their response at a fixed position over 1 s displays the periodic pattern illustrated in Fig. 1. However, modelled hippocampal cells have neither built-in spatial nor temporal properties. After learning, the response of modelled hippocampal cells is determined by the model dynamics (Eq. 7) where the connection weight matrix **A** is learnt during the virtual rat navigation. To investigate the temporal properties of learnt hippocampal place cells after learning, the response of learnt hippocampal place cells over 1 s at a given location is fitted to the following function:

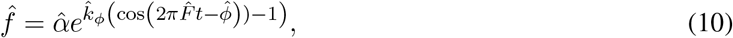

where 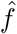 denotes the fitted response, 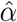 is the fitted amplitude that represents the spatial firing rate at this location, 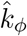 is the estimated parameter that controls the level of dependence on phase precession, 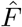 is the estimated frequency of the response, and 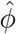 is the estimated phase of the response. The fitting routine is performed using the built-in MATLAB (version R2020a) function *lsqcurvefit*. The fitting error is defined as the square of the ratio between the fitting residual and original response.

### 2.9 Measuring the temporal properties of learnt hippocampal place cells

In order to quantitatively investigate the temporal properties of theta phase precession of learnt hippocampal place cells after learning, the following approach was used. First, for the learnt hippocampal place cells that meet the criteria of a place cell described earlier (Section 2.7.2), only cells whose entire place fields lie in the spatial environment are considered. Second, for each learnt hippocampal place cell, a virtual rat starts from the left side of the place field with an initial running rightwards direction and then a curved trajectory is generated according to Eq. 1. Third, the response over 1 s at each position along the trajectory is generated by the model and then fitted using Eq. 10. Finally, the entry phase, exit phase, and correlation coefficient between firing phases and normalised pdcd are obtained. For a learnt hippocampal place cell with theta phase precession, entry phase should be large and close to 360°, exit phase should be small and close to 0°, and the correlation coefficient between firing phases and normalised pdcd should be negative and close to −1.

## 3 Results

### 3.1 Scenario 1: the spatio-temporal properties of hippocampal place cells can be inherited from MEC grid cells via plasticity

The results presented here are those from Scenario 1, namely the simulation in which only MEC spatio-temporal grid cells provide input for the modelled hippocampal cells. The MEC spatio-temporal grid cell to hippocampal cell connectivity was learnt using non-negative sparse coding over 3600 s of virtual rat navigation over the 1 m × 1 m environment, as described in Section 2.6. Through this learning, the learnt hippocampal place cells learn to pool different MEC spatio-temporal grid cells in such a way that they become selective to specific locations in the spatial environment. The receptive fields of the 94 out of 100 modelled hippocampal cells that meet the criteria for place cells described in Section 2.7.2 are plotted in Fig. 3A. These learnt hippocampal place cells have a dominant firing location and their firing locations differ from each other. The centres of each of these 94 place fields are displayed in Fig. 3B, which are estimated using Eq. 9. This figure shows that the population of 94 learnt hippocampal place cells tile the entire spatial environment in a fairly uniform fashion. Compared with our previous work (Fig. 3 in Lian and Burkitt (2021)), the tiling of hippocampal place cells here is less uniform due to the lack of uniformity in the input spatial locations: the spatial locations in Lian and Burkitt (2021) are randomly sampled from a uniform distribution while the spatial locations in this paper are generated by a rat running trajectory described in Eq. 1.

**Figure 3:**
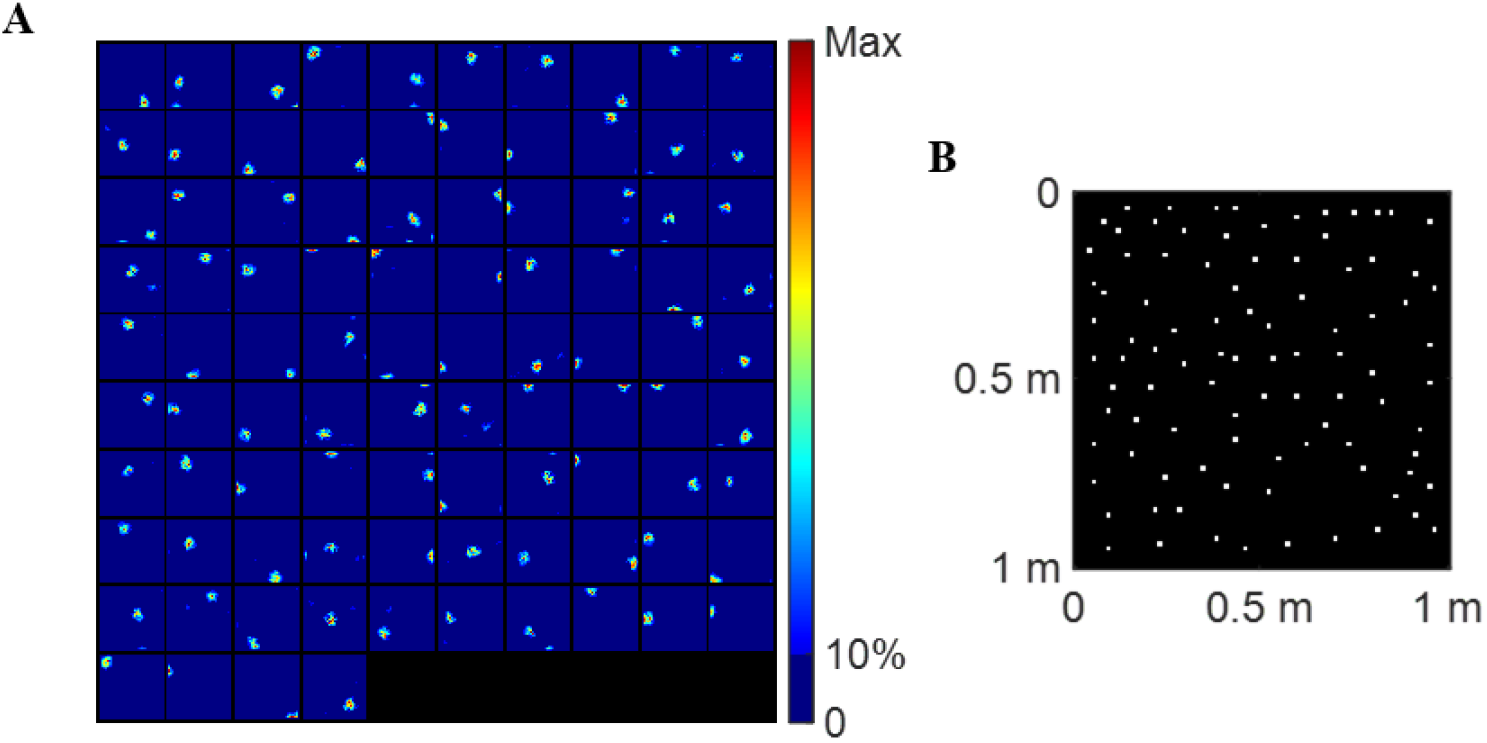
Scenario 1: learnt hippocampal place cells display spatial properties. (A) Place fields of 94 (out of 100) learnt hippocampal place cells that satisfied the three criteria for a place cell. Each block represents the spatial receptive field (rate map) of a cell in a 1m × 1m environment. (B) Centres of these 94 learnt hippocampal place cells plotted together in the 1m × 1m environment. Values of parameters used in the simulation are given in Table 1.

Furthermore, these learnt hippocampal place cells also display the temporal property of theta phase precession. This is illustrated in the response of a learnt hippocampal place cell over 0.5 s shown in Fig. 4.

**Figure 4:**
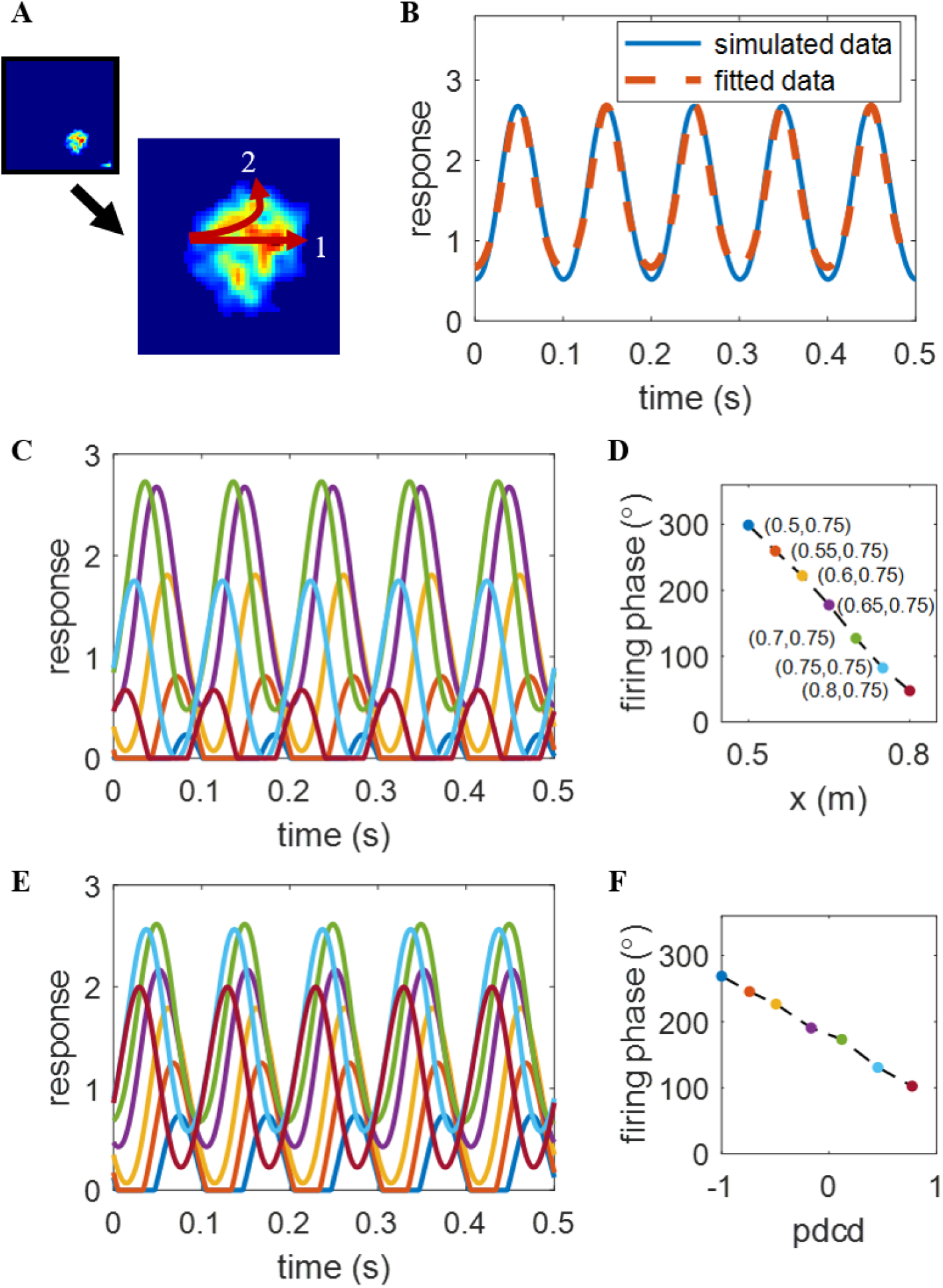
Scenario 1: one example of learnt hippocampal place cell’s temporal response properties. (A) Rate map of an example of learnt place cells whose centre is at (0.65, 0.75). (B) The response (solid line) over 0.5 s at the centre of the place field and the fitted response (dashed line) by Eq. 10. (C) The curves in different colours represent responses of this learnt hippocampal place cell over 0.5 s at different locations (coordinates found in plot D with the same colour coding) along the straight trajectory ‘1’ in plot A. (D) Estimated firing phase, 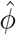, vs. position. Only *x* varies because the virtual rat moves from left to right. (E) The curves in different colours represent responses of this learnt hippocampal place cell over 0.5 s at different locations (normalised pdcd found in plot F with the same colour coding) along the curved trajectory ‘2’ in plot A. (F) Estimated firing phase, 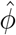, vs. normalised pdcd. Values of parameters used in the simulation can be found in Table 1.

Fig. 4B shows the response of one learnt hippocampal place cell over 0.5 s at position (0.65, 0.75), which is the centre of the place field shown in Fig 4A. The solid line in Fig 4B represents the response of this example place cell over 0.5 s and displays a periodic pattern. After fitting the response of the learnt place cell to the spatio-temporal function described by Eq. 10, the fitted response is plotted in the dashed line. The response of this learnt place cell is well fitted by Eq. 10 and the fitting error is small (0.35%). The fitted parameters are 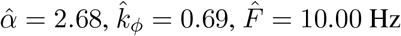 and 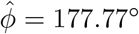.

The response of the same example place cells over 0.5 s as the animals runs from left to right of the place field (through the field centre) along the straight trajectory ‘1’ in Fig. 4A is shown in Fig. 4C. As the virtual rat is at position (0.5, 0.75), the amplitude of the response curve is below 1 and the firing phase (indicated by the peak location) is close to the end of each cycle (i.e., late phase). As the virtual rat continues running to the right, the amplitude of the response curve increases and then decreases, while the firing phase continues to shift forward to the beginning of each cycle. After fitting the response curve by Eq. 10, the relationship between firing phase and position is shown in Fig. 4D, which shows a clear reverse correlation; namely that the firing phase moves from late phase to early phase as the virtual rat crosses the place field. The entry phase and exit phase are 303.9° and 28.7°, respectively. The correlation coefficient between firing phases and positions is −0.998. Therefore, as the virtual rat runs from the left to the right of the place field, this place cell response displays theta phase precession. In other words, after learning, the resulting hippocampal place cells not only display spatial place fields in their responses but also display phase precession.

The learnt place cell also displays theta phase precession when the virtual rat crosses the place field along the curved trajectory ‘2’ in Fig. 4A, as illustrated in Fig. 4E & F. Since in this case both x and y values of the position change, we plot the firing phase relative to the normalised pdcd (similar to Jeewajee et al. (2014)) instead of positions. The amplitude of the response curve is small when the virtual rat enters the place field, increases to the peak, and then decreases as the virtual rat leaves the place field. The firing phase is observed to shift from late phase to early phase; i.e., displaying theta phase precession, similar to the straight trajectory but with a somewhat narrow range of phases. The entry phase is 268.8°, exit phase is 102.3°, and the correlation coefficient between firing phases and normalised pdcd is −0.998.

Consequently the example place cell in Fig. 4 learns both spatial and temporal properties after training. These spatio-temporal properties for the learnt hippocampal place cells are not inherent properties of the place cells, but arise entirely due to the way in which the model learns a place-specific tuning and theta phase precession from the spatio-temporal information provided by MEC grid cells.

Among the 94 (out of 100) learnt hippocampal place cells, 58 place cells have their entire place fields inside the spatial environment. For each place cell, a random curved trajectory is generated that crosses the place field and the neuronal response is computed at different locations along the trajectory (Section 2.9). The population statistics of the temporal properties are shown in Fig. 5. Fig. 5A & B show that the firing phase of these cells is at a late phase when the virtual rat enters the place field and at an early phase when the virtual rat leaves the place field. The histogram of correlation coefficients between firing phase and normalised pdcd (Fig. 4C) and scatter plot of firing phase and normalised pdcd (Fig. 4D) demonstrate strong theta phase precession. Recall that MEC grid cells are constructed here to have this built-in phase precession property (characterised by Eqs. 4 and 6). For each grid field of MEC grid cell, the firing phase and normalised pdcd along any trajectory are constructed to have a correlation coefficient of −1 due to the linear relationship between the firing phase and pdcd (Eq. 6). Both the spatial and temporal response properties of hippocampal place cells are entirely learnt from pooling the input they receive from MEC grid cells. Furthermore, the fact that the correlation coefficients in Fig. 5 are very close to −1 indicates that theta phase precession is well preserved in the learnt hippocampal place cells.

**Figure 5:**
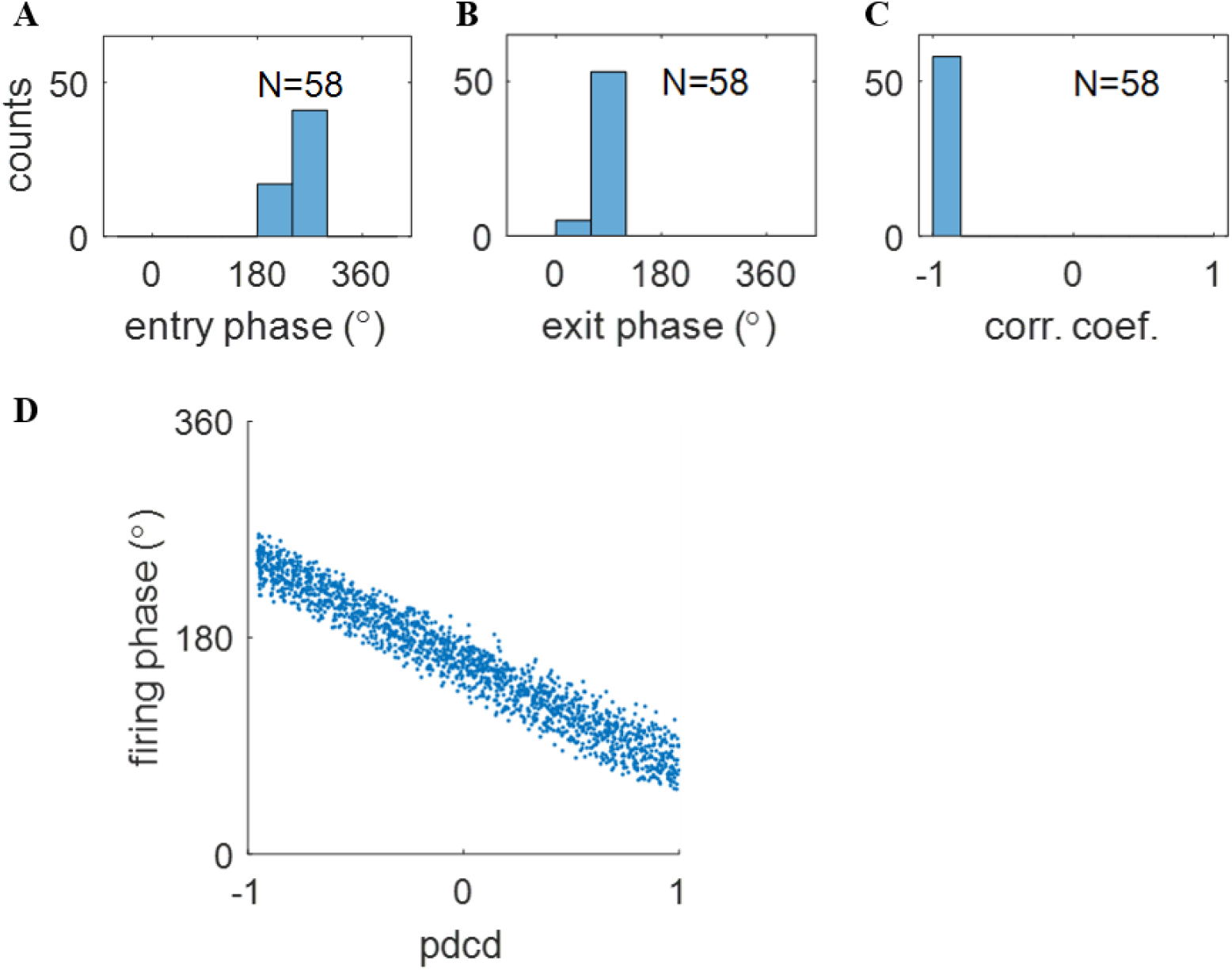
Scenario 1: the population of learnt hippocampal place cells displays temporal properties. Histograms of (A) the entry phases, (B) the exit phase, and (C) the correlation coefficient show that learnt hippocampal place cells have strong theta phase precession. (D) Scatter plot of firing phase, 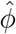, vs. pdcd, clearly showing theta phase precession of learnt hippocampal population of cells. Values of parameters used in the simulation are given in Table 1.

### 3.2 Scenario 2: the spatio-temporal properties of hippocampal place cells can be learnt when both MEC grid cells and EC weakly spatial cells serve as the EC input to the hippocampus

In this section, we show the results of training with Scenario 2, namely where both MEC spatio-temporal grid cells and EC weakly spatial cells provide input to the modelled hippocampal cells. The results show that both MEC grid cells and EC weakly spatial cells provide spatial information for the hippocampus, so that a spatial hippocampal map can be retrieved from upstream input, while MEC grid cells provide temporal information that results in theta phase precession also being inherited from the upstream neural population.

A demonstration of how EC weakly spatial cells and MEC grid cells contribute to the spatial and temporal properties of hippocampal place cells is given in Fig. 6. After learning, the recovered rate maps show place-field like receptive fields. As seen from Fig. 6A, different hippocampal place cells learn different place fields and the hippocampal population tiles the entire environment, similar to Scenario 1 in which only MEC grid cells provide input to the hippocampus (Fig. 3). An example of the temporal response properties of a place cell can be seen from the relationship between firing phase and normalised pdcd within the place field in Fig. 6B, which displays characteristic theta phase precession: as the virtual rat moves along a random generated curved trajectory (Section 2.9), the firing phase shifts smoothly from a late phase to an early phase. Fig. 6C & D displays the population statistics for this phase precession, similar to Fig. 5, indicating that learnt hippocampal place cells display strong phase precession.

**Figure 6:**
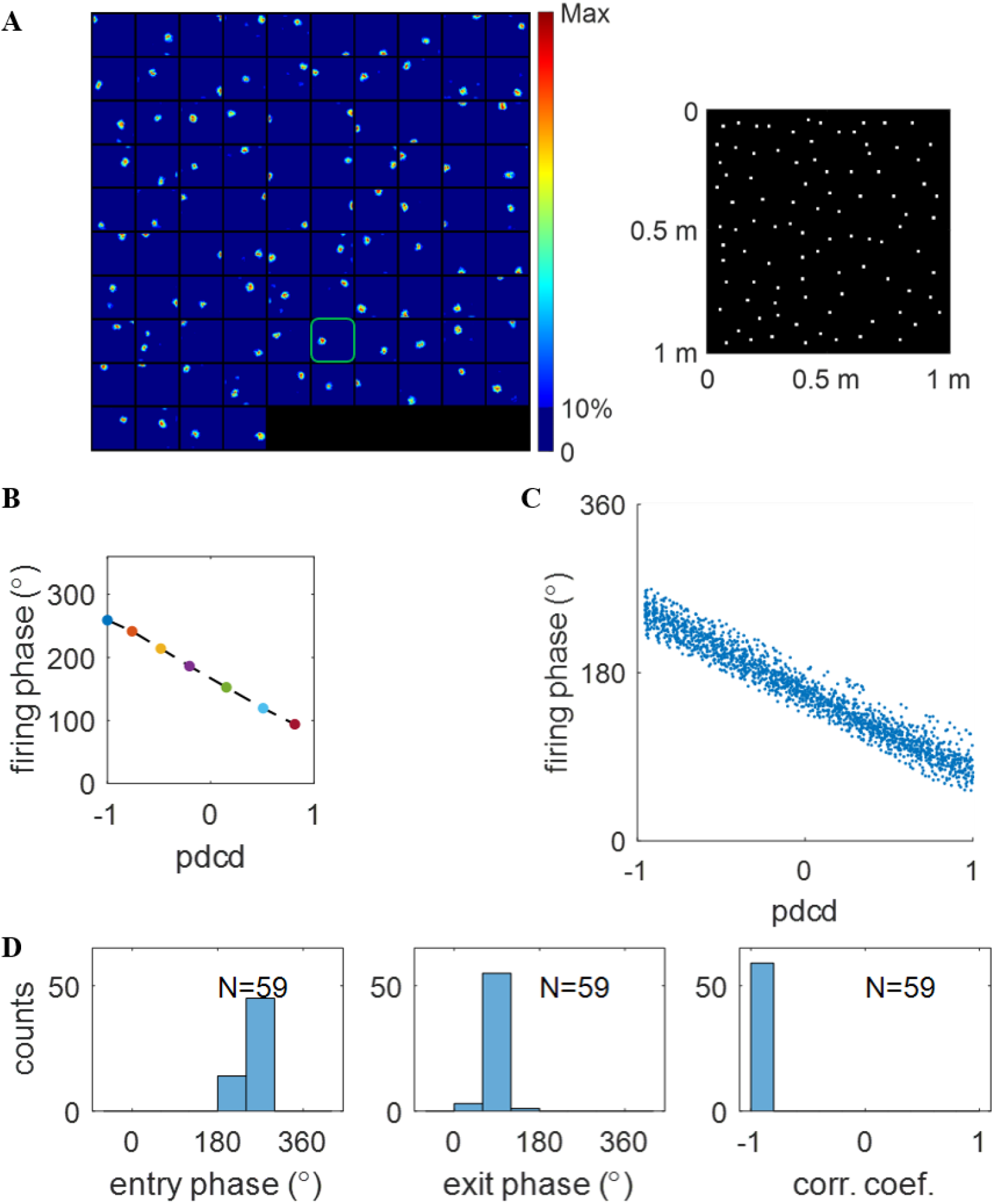
Scenario 2: learnt hippocampal place cells display spatio-temporal properties. (A) Spatial properties of learnt hippocampal place cells. Left: the place fields of learnt hippocampal place cells; Right: place field centres plotted in the same environment. (B) Firing phase, 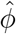, vs. normalised pdcd along a curved trajectory that crosses the place field, which illustrates theta phase precession for one learnt hippocampal place cell highlighted in plot A. (C) The scatter plot of firing phase, 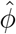, vs. pdcd, clearly showing theta phase precession of learnt hippocampal population. (D) The population of learnt hippocampal place cells displays temporal properties. Histograms of entry phase (Left), exit phase (Middle), and correlation coefficient (Right) indicate a strong phase precession for the population of learnt hippocampal place cells. Values of parameters used in the simulation are given in Table 1.

### 3.3 Scenario 3: spatial properties of hippocampal place cells are maintained after the inactivation of MEC grid cells

After the learning process in Scenario 2, MEC spatio-temporal grid cells in the model were inactivated; i.e., only the remaining EC weakly spatial cells provide spatial information for hippocampal place cells and there is no input (temporal or spatio-temporal) from MEC grid cells. Recall that there is no learning in Scenario 3, so the spatial tuning of hippocampal place cells depends on the connectivity between EC weakly spatial cells and hippocampal place cells that was learnt in Scenario 2.

However, the learnt connection between EC weakly spatial cells and place cells is sufficient to maintain the place field of hippocampal place cells after MEC inactivation. After recovering the receptive fields of modelled hippocampal cells (see Section 2.7), 87 out of 100 modelled hippocampal cells were found to meet the criteria of a place cell. The place fields, together with their field centres, are plotted in Fig. 7A. Comparison with Fig. 6A shows that the place fields still maintain their firing locations and the population still tiles the entire spatial environment. This indicates that the learnt connection from EC weakly spatial cells to place cells during free running provides sufficient spatial information, such that stable place fields will not be lost even though MEC grid cells are inactivated in this Scenario.

**Figure 7:**
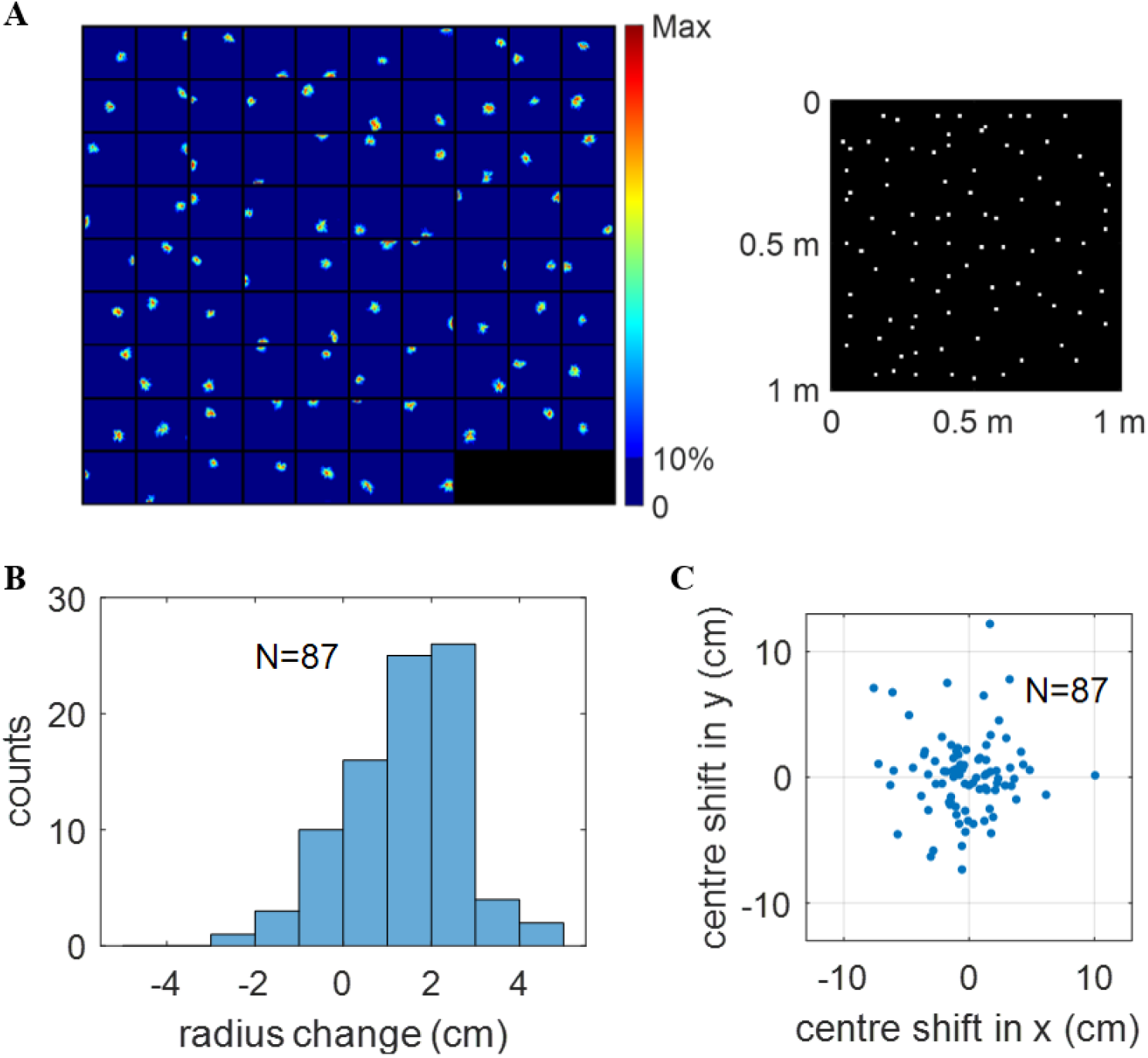
Scenario 3: learnt hippocampal place cells maintain spatial firing after MEC inactivation. (A) The place fields of 87 hippocampal place cells (Left) and their centres plotted in the same environment (Right). (B) Histogram of radius change of place fields after MEC inactivation. The radius of most place fields increases after MEC inactivation. The mean radii before and after MEC inactivation are 8.94 cm and 10.34 cm, respectively. (C) Scatter plot of the place field centre shift after MEC inactivation.

There are 94 place cells before inactivating MEC grid cells (Fig. 6A, Scenario 2) and 87 place cells after the inactivation (Fig. 7A, Scenario 3). We find that all these 87 place cells that pass the criteria for place cells after MEC inactivation (Scenario 3) come from the population of 94 place cells before MEC inactivation (Scenario 2). After inactivating MEC grid cells, Fig. 7B shows that, in general, the size of the place fields increases. The mean radii before and after MEC inactivation are 8.94 cm and 10.34 cm, respectively. Furthermore, although most place cells maintain stable place fields after the inactivation, their field centres shift randomly within a small range, as seen in Fig. 7C. These results also suggest that the place coding of the hippocampus becomes somewhat less accurate after MEC inactivation.

This result is consistent with the experimental study by Schlesiger et al. (2015) who found that after MEC lesions in rats, theta phase precession of hippocampal CA1 cells was significantly disrupted, although stable spatial firing was maintained. Therefore, during the navigation of the virtual rat in Scenario 2, the plasticity of the model allows hippocampal place cells to pool the input from upstream neurons, namely the EC weakly spatial cells or MEC grid cells, enabling both spatial and temporal properties to be learnt.

Combining results in this paper and our previous work (Lian and Burkitt, 2021), the contribution of the EC to the firing of hippocampal place cells can be described as follows: both EC weakly spatial cells and cells with a clear structure (such as MEC grid cells) provide spatial information for the hippocampus to learn an efficient hippocampal place map, while the temporal response properties involving theta phase precession of hippocampal place cells are inherited from MEC grid cells via learning during navigation.

## 4 Discussion

In this study, a model based on sparse coding is built that demonstrates that the spatial and temporal properties of hippocampal place cells can be learnt simultaneously via plasticity as the virtual rat freely explores an open environment. In the training phase, a virtual rat runs for 3600 s and the connectivity weight matrix, **A**, is learnt during this exploration period. After learning, **A** is kept fixed and another running trajectory of 1200 s is used to recover the place fields of learnt place cells. In addition, responses over 1 s at different positions across the place field are used to investigate the firing phases when the virtual rat traverses the place field. Similar to the study of Jeewajee et al. (2014), pdcd is used as the measurement of position when the rat is running along a curved trajectory. Our results show that the learnt hippocampal place cells are selective to a single firing location and display theta phase precession, even though the learnt hippocampal place cells have no pre-built spatial and temporal properties. Furthermore, the model shows that the loss of MEC grid cells causes the loss of the temporal response properties of hippocampal place cells, but the spatial properties of hippocampal place cells are maintained by other upstream cells, such as EC weakly spatial cells, that provide spatial information for the hippocampus. This model demonstrates how the spatio-temporal properties of hippocampal place cells can be learnt.

### 4.1 The entorhinal-hippocampal loop

Early computational models of place cells were mostly based upon a feedforward structure, where cells in the EC provide spatial input to the hippocampus (Solstad et al., 2006; Franzius et al., 2007b,a; de Almeida et al., 2009). Some more recent studies have adopted a loop network structure in which cells in the EC project to the hippocampus and also receive feedback from it (Rennó-Costa and Tort, 2017; Li et al., 2020; Agmon and Burak, 2020). Models incorporating this loop network structure can explain more experiment observations, especially on how hippocampal place cells affect the firing of MEC grid cells. Since the hippocampus receives information from different brain areas via the EC, place cells in the hippocampus can still maintain some properties even when MEC grid cells are inactivated. On the other hand, hippocampal place cells also affect the grid pattern of MEC grid cells (Bonnevie et al., 2013; Almog et al., 2019). Therefore, the feedback from the hippocampus to the EC can modify some properties of EC cells, which can only be investigated using the loop network structure. However, the results presented here do not rule out the possibility of implementing the model in a loop network structure because sparse coding can be implemented in a feedforward-feedback loop (Lian et al., 2019).

### 4.2 Evidence against feedforward grid-to-place models

Recent research has provided experimental evidence that place fields emerge earlier in development than MEC grid cells (Langston et al., 2010; Wills et al., 2010). There is also experimental evidence for the maintenance of stable place fields after the inactivation of MEC grid cells (Brandon et al., 2014; Schlesiger et al., 2015; Hales et al., 2014). These results put into question the feedforward nature of grid-to-place cell models. However, unlike typical grid-to-place models, our model takes input from different types of cells in the EC. The results presented in this paper, together with our previous work (Lian and Burkitt, 2021), provides a unifying framework that is able to explain this experimental evidence while supporting a hierarchical structure of connectivity from the EC to the hippocampus in which theta phase precession of hippocampal place cells is essentially inherited from MEC gird cells via learning.

Bonnevie et al. (2013) showed that the inactivation of place cells will cause the loss of MEC grid cells. However, Almog et al. (2019) re-analyzed the same data and found that grid cells maintain synchrony even though grid tuning is lost. In addition, a large number of grid cells actually maintain spatial tuning after the reactivation of place cells (Fig. 1D in Almog et al. (2019)). Therefore, though hippocampal place cells affect the spatial firing of some MEC grid cells, this does not stand against the importance of the feedforward connection from EC to the hippocampus. A future model that can capture the contribution of the EC to the hippocampus as well as the effect the hippocampus has on the EC is needed.

### 4.3 Future work

While this study offers a picture of how the EC contributes to the spatial and temporal properties of hippocampal place cell, there remain a number of interesting outstanding questions to be addressed. First, the question of how the feedback from the hippocampus to the EC affects properties of EC cells remains unclear. Second, the model proposed here demonstrates how sparse coding can learn spatio-temporal properties of hippocampal place cells, but whether the principle of sparse coding can be used to explain other aspects of hippocampal function, such as those involving memory consolidation, remains unclear. In addition, the spatio-temporal model of MEC grid cells is assumed here, and the question of how this MEC spatio-temporal grid cell structure originates is a topic of ongoing research. Moreover, the model presented here uses rate-based neurons: a spiking implementation together with spike-timing dependent plasticity of this model is left for future research.

## Acknowledgements

This work received funding from the Australian Government, via grant AUS-MURIB000001 associated with ONR MURI grant N00014-19-1-2571. We thank Michael E. Hasselmo for helpful comments on the manuscript.

